# Refining the domain architecture model of the replication origin firing factor Treslin/TICRR

**DOI:** 10.1101/2021.07.24.453613

**Authors:** Pedro Ferreira, Luis Sanchez-Pulido, Anika Marko, Chris P Ponting, Dominik Boos

## Abstract

Faithful genome duplication requires appropriately controlled replication origin firing. The metazoan Treslin/TICRR origin firing factor and its yeast orthologue Sld3 are regulation hubs of origin firing. They share the Sld3-Treslin domain (STD) and the adjacent TopBP1/Dpb11 interaction domain (TDIN). We report a revised domain architecture model of Treslin/TICRR. Complementary protein sequence analyses uncovered Ku70-homologous β-barrel folds in the Treslin/TICRR middle domain (M domain) and in Sld3. Thus, the Sld3-homologous Treslin/TICRR core comprises its three central domains, M domain, STD and TDIN. This Sld3-core is flanked by non-conserved terminal domains, the CIT (conserved in Treslins) and the C-terminus. We also identified Ku70-like β-barrels in MTBP and Sld7. Our binding experiments showed that the Treslin β-barrel mediates interaction with the MTBP β-barrel, reminiscent of the homotypic Ku70-Ku80 dimerization. This binding mode is conserved in the Sld3-Sld7 dimer. We used Treslin/TICRR domain mutants to show that all Sld3-core domains and the non-conserved terminal domains fulfil important functions during origin firing in human cells. Thus, metazoa-specific and widely conserved molecular processes cooperate during origin firing in metazoa.

## Introduction

Accurate and complete DNA replication guarantees faithful genetic inheritance. It requires complex regulation of replication origin firing to ensure 1) efficient firing to avoid non-replicated gaps, and 2) appropriately controlled firing in space and time to facilitate the metazoan genome replication program and coordinate replication with other chromatin processes like transcription (Berezney et al 2000, Boos & Ferreira 2019, Dileep et al 2015, Helmrich et al 2013, Petryk et al 2016, Ryba et al 2010).

Replication initiation is a two-step process in eukaryotes. The first step, origin licensing, in G1 phase is the formation of pre-replicative complex (pre-RC), the loading of the Mcm2-7 replicative helicase onto double-stranded DNA (Evrin et al 2009, Remus et al 2009). In pre-RCs, the Mcm2-7 complex does not have helicase activity to avoid premature DNA unwinding in G1. The second step is origin firing, the conversion of pre-RCs into two bidirectional replisomes. Firing occurs S phase-specifically due to its dependency on S phase cyclin-dependent kinases (S-CDK) and Dbf4-dependent kinase (DDK), whose activities increase at the G1-S transition. During firing, pre-RCs are first remodelled into pre-initiation complexes (pre-ICs) (Miyazawa-Onami et al 2017, Yeeles et al 2015, Zou & Stillman 1998) that then mature into the active Cdc45-Mcm2-7-GINS-DNA polymerase epsilon (CMGE) helicase (Abid Ali et al 2017, Douglas et al 2018, Ilves et al 2010, Langston et al 2014). DNA synthesis requires assembly of additional replisome factors and primer synthesis (Yeeles et al 2017).

The main regulation step of origin firing is pre-IC formation. In yeast, a dimer of Sld3 and Sld7 (orthologues of metazoan Treslin/TICRR and MTBP (Kohler et al 2019, Kumagai & Dunphy 2017, Sanchez-Pulido et al 2010)), binds pre-RCs dependently on pre-RC phosphorylation by DDK (Deegan et al 2016, Heller et al 2011). Sld3 recruits Cdc45 via its central STD domain (Itou et al 2014, Kamimura et al 2001) (Fig. 1). Sld3 utilizes its TDIN region to bind to Dpb11 (TopBP1/Cut5/Mus101 in higher eukaryotes) in an interaction that depends on phosphorylation at two CDK sites in the TDIN (Boos et al 2011, Zegerman & Diffley 2007). Dpb11 also binds CDK-phosphorylated Sld2 (RecQL4 in higher eukaryotes). Dpb11 and Sld2 form the pre-loading complex together with GINS and DNA polymerase epsilon (Muramatsu et al 2010). The resulting intermediate structure is called pre-IC. Then, Sld3, Dpb11 and Sld2 dissociate and the CMGE helicase forms.

**Figure 1.**
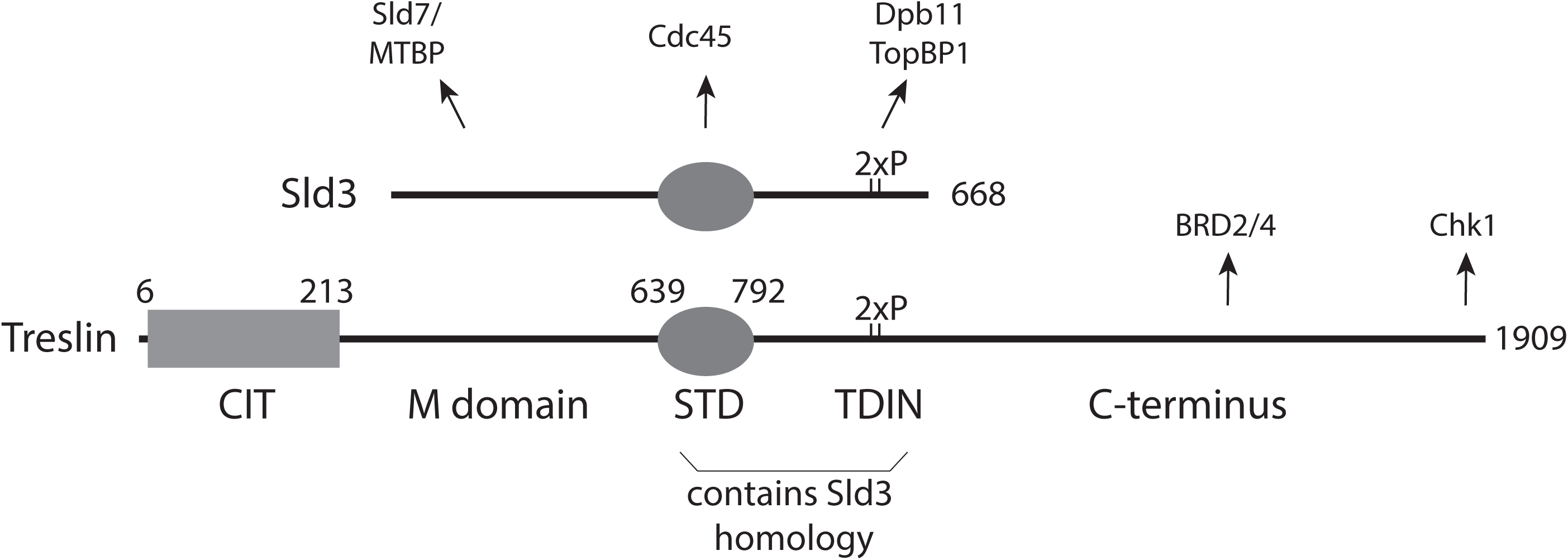
Treslin/TICRR domain structure. CIT: Conserved in Treslins; M: middle domain; STD: Sld3-Treslin domain; TDIN: TopBP1/Dpb11 interaction domain. Numbers indicate amino acid position in human Treslin/TICRR or budding yeast Sld3. Arrows point to interacting proteins. 2xP; two phospho-serines.

In addition to cell cycle kinases, the DNA damage checkpoint also controls origin firing at the pre-IC step. Checkpoint kinase phosphorylation of Sld3 and Dbf4 inhibit pre-IC formation to avoid mutations through replicating damaged templates (Duch et al 2011, Lopez-Mosqueda et al 2010, Zegerman & Diffley 2010). Recently, it has become clear that more subtle regulation of pre-IC factor activity and levels are critical for faithful genome duplication in yeast (Mantiero et al 2011, Reusswig et al 2016, Tanaka & Araki 2011, Tanaka et al 2011).

Many fundamental processes of yeast origin firing are conserved in vertebrates. All yeast origin firing factors have orthologues in higher eukaryotes (Kohler et al 2019). In addition, cell cycle regulation by CDK through Treslin/Sld3 binding to TopBP1/Dpb11 and also firing inhibition upon DNA damage through suppression of the Treslin/Sld3-TopBP1/Dpb11 interaction are both conserved (Boos et al 2011, Guo et al 2015, Kumagai et al 2010; 2011, Mu et al 2017, Sansam et al 2010).

However, several protein domains of TopBP1, MTBP, Treslin/TICRR and RecQL4 do not have counterparts in yeasts (Kohler et al 2019, Makiniemi et al 2001, Sanchez-Pulido et al 2010, Zegerman 2015). This suggests that, despite the described conservation, metazoa and fungi have evolved specific origin firing processes. Whilst it has been shown that some higher eukaryote-specific domains of MTBP and TopBP1 are required for efficient DNA synthesis (Kohler et al 2019, Kumagai et al 2010), the situation for Treslin/TICRR remains less clear. Characterisation of the protein domains that are specific to higher eukaryotes is essential for defining how origin firing processes in these cells diverge from the established yeast model.

The two central STD and TDIN domains of Treslin/TICRR show sequence-based evidence for homology with Sld3 (Fig. 1) (Boos et al 2011, Itou et al 2014, Sanchez-Pulido et al 2010). The molecular functions of the STD of Treslin/TICRR and whether this region is essential for replication remain unknown. Its homology with Sld3 suggests that it might support origin firing through interaction with Cdc45 (Itou et al 2014). The TDIN of Treslin/TICRR is a conserved region containing two CDK phosphorylation sites for TopBP1 binding (Boos et al 2011, Kumagai et al 2011). Like the Sld3-TDIN the Treslin/TICRR-TDIN forms a direct binding surface for BRCA1 C-terminal repeat domains (BRCT) in TopBP1/Dpb11 (Boos et al 2011, Kumagai et al 2011, Zegerman & Diffley 2007).

The Treslin/TICRR domains N- and C-terminal of STD and TDIN (Fig. 1) have not been shown to be conserved with Sld3. The M domain shares the ability to bind to MTBP/Sld7 with the N-terminal region of Sld3, and it is required for replication in human cells (Itou et al 2015, Kohler et al 2019). It came as a surprise that sequence conservation with Sld3 was not detected for the Treslin/TICRR M domain, because the interacting regions in MTBP and Sld7, respectively, show homology via remote but statistically significant sequence similarity (Kohler et al 2019). The region C-terminal of the TDIN is present in many metazoans, but is absent from yeast and plants (Sanchez-Pulido et al 2010). Sequence analysis predicts that this Treslin/TICRR C-terminal region is largely unstructured, with well-conserved stretches of amino acids and more divergent regions alternating. This region binds Chk1 and BRD2/4 (Fig. 1), but these activities are not essential for DNA synthesis in cultured human cells (Guo et al 2015, Sansam et al 2018). The N-terminal CIT is conserved in both metazoans and plants, but not present in fungi (Sanchez-Pulido et al 2010). Whether the CIT functions in replication is unknown.

We here define the essential Sld3-like core of Treslin/TICRR as the three M, STD and TDIN domains, flanked by higher eukaryote-specific terminal domains. Moreover, we characterise structurally and functionally the M domain and the higher eukaryote-specific terminal regions.

## Results

### The M domain, the STD and the TDIN domain constitute the essential core of Treslin/TICRR

We first sought to better define the essential core domains of Treslin/TICRR for replication. Mutations of Treslin/TICRR previously showed that the MTBP/Sld7-binding M domain and the TopBP1/Dpb11-binding TDIN perform essential functions during origin firing in human cells (Boos et al 2011, Boos et al 2013, Kumagai & Dunphy 2017). In contrast, the requirement of the Sld3-homologous STD for replication had not previously been addressed in higher eukaryotes. To test this, we used incorporation of the nucleotide analogue 5-bromo-2’-deoxyuridine (BrdU) into nascent DNA of cultured human cells in an established RNAi- replacement system (Boos et al 2011, Boos et al 2013). U2OS cell clones stably expressing siRNA-resistant Treslin/TICRR wild type (WT) or STD-deletion mutants (ΔSTD, amino acids 717-792 deleted) to similar levels (Fig. 2A) were treated with control siRNA (siCtr) or Treslin/TICRR siRNA (siTreslin). 72 h after transfection cells were pulse-labelled with BrdU, stained with anti-BrdU-FITC and propidium iodide (PI), and analysed by flow cytometry. Parental U2OS cells and control cell lines expressing the inactive non-TopBP1 interacting CDK site mutant Treslin/TICRR-2PM showed severely reduced BrdU incorporation levels compared to siCtr-treated cells (Fig 2B). Whilst Treslin/TICRR-WT rescued BrdU incorporation, three independent clones expressing Treslin/TICRR-ΔSTD (clones 11, 17 and 21) showed strong defects in supporting replication (Fig 2B). Quantification of replicates (Fig. 2C) confirmed these observations and proved reproducibility of our BrdU quantification method (Boos et al 2013, Ferreira et al 2021, Kohler et al 2019). Treslin/TICRR-ΔSTD clone 21 rescued replication somewhat better (50 % replication) than clones 11 and 17 (approximately 30 % replication; 2PM and no-transgene controls about 30%), exemplifying our observation that individual clones expressing the same transgene showed some variability that probably arise through clonal selection. We then tested if specifically the origin firing step of replication is impaired in Treslin-ΔSTD cells by analysing origin licensing and replisome formation on chromatin. Western blotting of chromatin fractions using anti-Mcm2 antibodies showed that replication origin licensing occurred normally in the G1 phase (4 h after Nocodazole release) in Treslin/TICRR-ΔSTD cells. In contrast, origin firing did not occur in the absence of the STD domain as indicated by severely reduced S phase-specific (12 h) Cdc45 and PCNA loading onto chromatin (Fig 2D). The loss of replication activity is not a consequence of a delay in S phase entry, because Cyclin A accumulated normally in Treslin-ΔSTD cells 12 h after release (Fig 2D). STD deletion neither lead to gross misfolding of Treslin/TICRR nor affected the described activities of the neighbouring M and TDIN domains, because Treslin-ΔSTD immunoprecipitated MTBP (Boos et al 2013) and TopBP1 (Fig. S1) normally. We concluded from these RNAi-rescue experiments that deleting the STD severely compromises replication origin firing in U2OS cells. Thus, the STD is part of the essential set of core domains of Treslin/TICRR, together with the M and TDIN domains.

**Figure 2.**
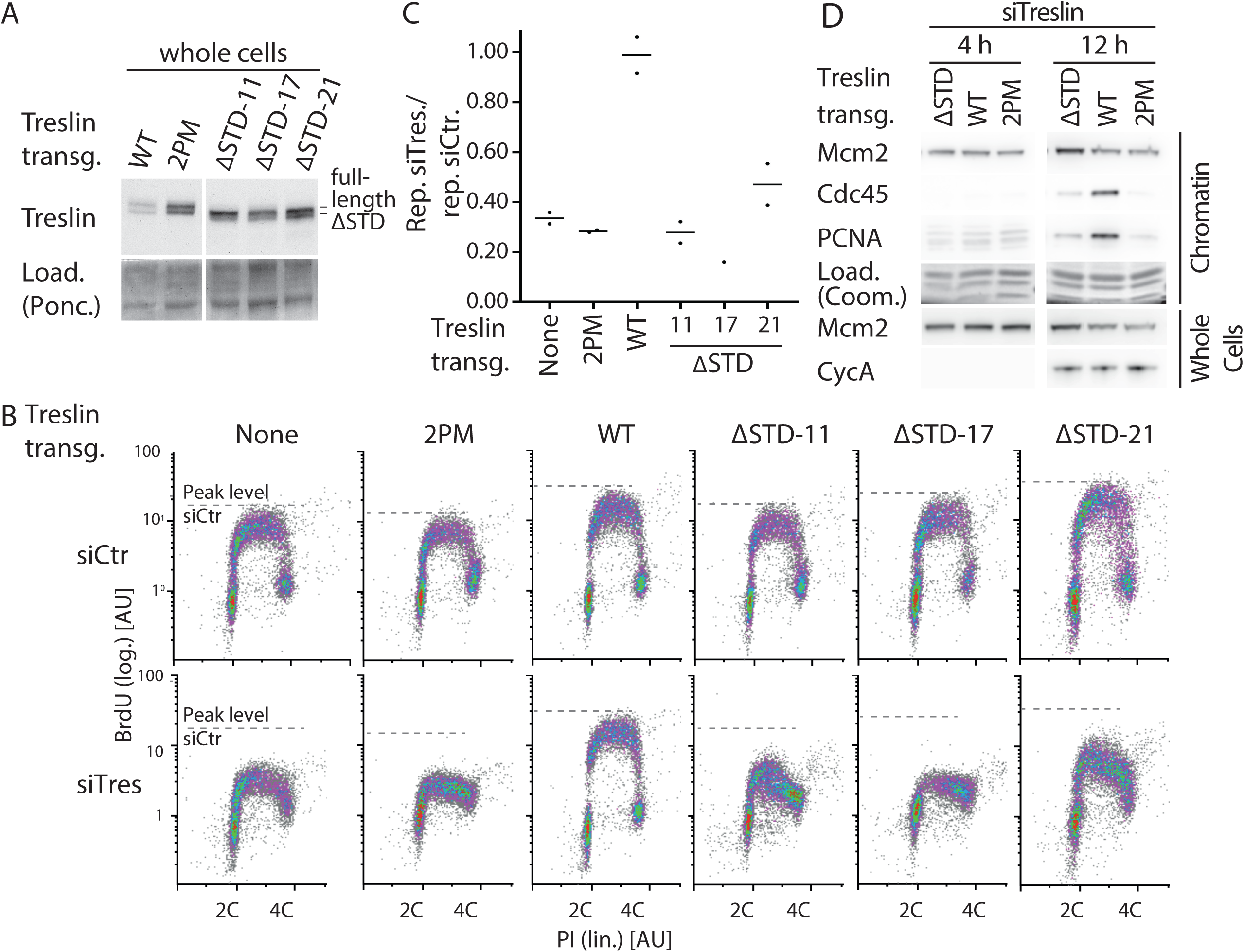
The STD domain of Treslin/TICRR is required for DNA replication in cultured human cells. **(A)** Whole cell lysates of stable U2OS cell lines carrying siRNA resistant transgenes of Treslin/TICRR-WT, Treslin/TICRR-2PM (threonine 969 and serine 1001 double alanine mutant that cannot interact with TopBP1(Boos et al 2011)) or three clones of Treslin/TICRR with a deletion of the STD (amino acids 717-792 deleted) were immunoblotted with rabbit anti-Treslin/TICRR (amino acids 1566-1909) antibodies. Ponceau (Ponc.) staining controlled for loading (Load.). **(B)** Cells described in A were treated with control or Treslin/TICRR siRNAs (siCtr/siTres) before analysis by flow cytometry detecting BrdU (5-bromo-2’-deoxyuridine; logarithmic (log.) scale) and PI (propidium iodide; linear (lin.) scale). Density plots are shown. Dashed lines indicating peak level of maximal BrdU incorporation in each cell line upon siCtr-treatment allow visual comparison with level upon siTres treatment. **(C)** Quantification of relative overall DNA replication in cells described in A based on flow cytometry experiments described in B. Averages of BrdU-replication signals of two (all lines except ΔSTD-17) or one (ΔSTD-17) experiments. Replication signals of siTreslin-treated cells were normalised to replication signals of the same cell line upon siCtr-treatments. **(D)** Stable U2OS cell lines expressing siTreslin-resistant Treslin/TICRR-ΔSTD, WT or 2PM were released from a double thymidine arrest before treatment with siTreslin and nocodazole. After nocodazole-release for 4 h or 12 h cells chromatin was isolated for immunoblotting with goat anti-Mcm2, rat anti-Cdc45 and mouse anti-PCNA antibodies. Whole cell lysates from the same samples were immunoblotted using mouse anti-cyclin A and goat anti-Mcm2 antibodies. For each antibody, crops are from the same immunoblot exposure. Coomassie (Coom.) staining of low molecular weight part including histones controlled for loading. Clone Treslin-ΔSTD -11 was used.

### Characterisation of the region N-terminal to the Treslin/TICRR-STD by protein sequence analysis

We then sought to better understand the region N-terminal to the STD of Treslin/TICRR, because it has no described sequence conservation with Sld3, but contains the M domain that has a conserved activity, the binding to MTBP/Sld7, and that is part of the essential Treslin/TICRR core. We used iterative protein sequence database searches seeking to characterize new domains and conserved regions in animal Treslin/TICRR. We anticipated that an increase in protein sequence and structure information and improved computational tools over the last ten years could complement our previous insights (Boos et al 2011, Sanchez-Pulido et al 2010).

First, we conducted a JackHMMER iterative search with amino acids 1-600 of the human Treslin/TICRR protein sequence (Eddy 1996, Finn et al 2011). This region is located N-terminal to the STD (amino acids 639-792) (Fig 1). As input for our domain hunting strategy, we first constructed a full-length multiple sequence alignment of Treslin/TICRR family members. During this process, we identified a N-terminal region of Treslin/TICRR (approximately amino acids 1-600) that is well-conserved across the animal kingdom, including earlier branching animals such as echinoderms, molluscs, annelids, and placozoans (*Trichoplax* sp.).

Then we used those sub-regions of this alignment that exhibited the highest levels of conservation as queries for profile-to-sequence (HMMer) and profile-to-profile comparisons (HHpred) (Eddy 1996, Finn et al 2011, Soding et al 2005). In particular, we undertook HHpred searches against the PDB70 profile database (Soding et al 2005), using the previously identified CIT region that is conserved between animal and plant Treslins (corresponding to residues 4 to 254 of human Treslin/TICRR) (Sanchez-Pulido et al 2010) (Fig 1). This search identified the Treslin/TICRR N-terminal domain as a von Willebrand factor type A (vWA) domain (also known as a Rossmann fold), with a highly significant match to the vWA domain of human complement factor B protein (PDB-ID: 3HRZ_D) (Janssen et al 2009) (*E*-value = 9.2 x 10^-3^; true positive probability of 97%) (Fig S2). In support of homology, the next most statistically significant matches were to more than ten additional members of the vWA domain superfamily, such as: human protein transport protein Sec23A (PDB-ID: 2NUT_A) (Mancias & Goldberg 2007), human Integrin β-8 (PDB-ID: 6DJP_B)(Cormier et al 2018), and yeast DNA repair protein Ku70 (PDB-ID: 5Y58_E) (Chen et al 2018). The secondary structure prediction for this region of Treslin/TICRR showed good agreement with the known secondary structure known of diverse members of the vWA superfamily (Jones 1999) (Fig S2).

The vWA domain consists of alternating β-strand and α-helix secondary structural elements, arranged in a central β-sheet (usually composed by a core of six β-strands) surrounded by the α-helices. The considerable functional diversity of vWA domain family members (Ponting et al 2000, Whittaker & Hynes 2002) disallows prediction of the potential functional role of the vWA domain using sequence comparison.

### The M domain of Treslin/TICRR contains a Ku70/Ku80-like dimerization domain and is similar to the N-terminal domain of Sld3

We next focused our bioinformatic analysis on the poorly characterised M domain. Although this Treslin/TICRR domain carries one of its core functions, one that is shared with Sld3 (the essential binding to MTBP/Sld7), its homology with Sld3 had not previously been demonstrated. HHpred searches of this region against the PDB70 profile database (Soding et al 2005) yielded statistically significant evidence of sequence similarity of Treslin/TICRR amino acids 341 to 426 to yeast Ku70 (PDB-ID: 5Y58_E) (Chen et al 2018) (*E*-value = 0.3; true positive probability of 88%) (Fig 3A). In further support of homology, the next most statistically significant matches were to three further members of the Ku family, namely yeast Ku80 (PDB-ID: 5Y58_F) (Chen et al 2018), human XRCC5 (PDB-ID: 1JEY_B), and human XRCC6 (X-ray repair cross-complementing protein 6) (PDB-ID: 1JEY_A) (Walker et al 2001).

**Figure 3.**
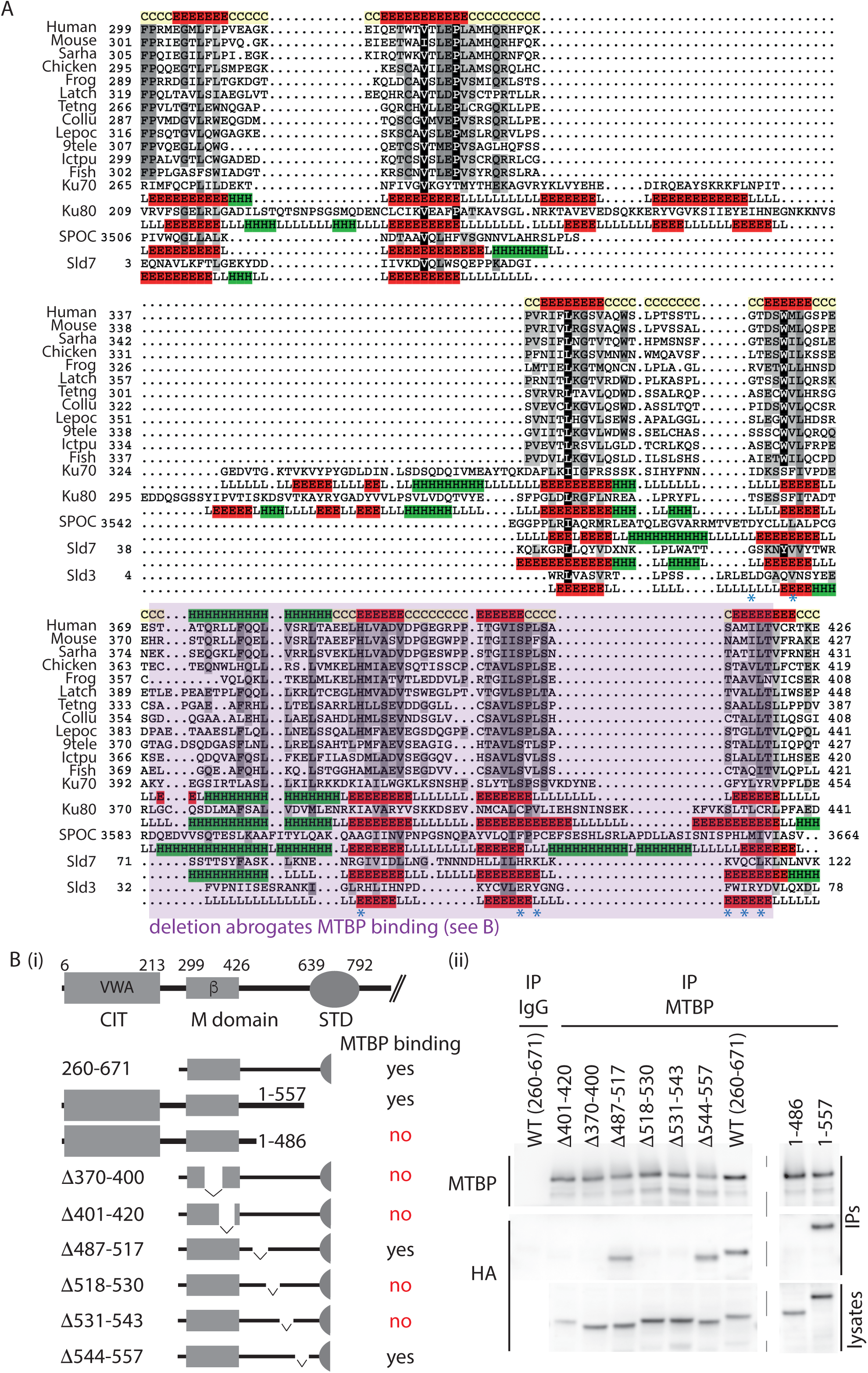
Treslin/TICRR, Sld3 and Sld7 contain a Ku70/80-like β-barrel that are required for Treslin/Sld3-MTBP/Sld7 dimerization. **(A)** Representative multiple sequence alignment of Ku70-like β-barrel domain in the Treslin/TICRR family. The alignment generated with the program T-Coffee (Notredame et al 2000) using default parameters and slightly refined manually. The final alignment was obtained using a combination of profile-to-profile comparisons (Soding et al 2005) and sequence alignments derived from structural super-positions of a selection of Ku70-like β-barrel domains whose tertiary structure is known (Holm & Sander 1995). The limits of the protein sequences included in the alignment are indicated by flanking residue positions. Secondary structure prediction using PsiPred (Jones 1999) was performed for the Treslin family, shown in the first lane; this prediction is consistent with the secondary structure of Ku70-like β-barrel domains, shown below each of the proteins with known structure (Ku70, PDB:5Y58E; Ku80, PDB:5Y58F; SPOC, PDB:1OW1A; Sld7, PDB:3X37B; Sld3, PDB:3X37A). Alpha-helices and β-strands are indicated by H and E, respectively. The alignment was presented with the program Belvu using a colouring scheme indicating the average BLOSUM62 scores (which are correlated with amino acid conservation) of each alignment column: black (>3), grey (between 3 and 1.5) and light grey (between 1.5 and 0.5) (Sonnhammer & Hollich 2005). Sequences are named according to their specie common name or abbreviation corresponding as follow to their UniProt identification and specie name (Wu et al 2006):Human, Q7Z2Z1_HUMAN, *Homo sapiens*; Mouse, Q8BQ33_MOUSE, *Mus musculus*; Sarha, G3WMD4_SARHA; *Sarcophilus harrisii*; Chicken, E1BU88_CHICK; *Gallus gallus*; Frog, D3IUT5_XENLA, *Xenopus laevis*; Latch, H3BCK8_LATCH, *Latimeria chalumnae*; Tetng, H3CYF8_TETNG, *Tetraodon nigroviridis*; Collu, A0A4U5UGV6_COLLU, *Collichthys lucidus*; Lepoc, W5ND48_LEPOC, *Lepisosteus oculatus*; 9tele, A0A3B3T1X9_9TELE, *Paramormyrops kingsleyae*; Ictpu, A0A2D0SG01_ICTPU, *Ictalurus punctatus*; Fish, Q6DRL4_DANRE, *Danio rerio.* Blue asterisks: amino acid positions in Sld3 that mediate Sld7 interaction (Itou et al 2015) **(B)** Schematic representation of Treslin/TICRR mutants (i) used for interaction studies (ii). For (ii), the indicated N-terminally 3HA-tagged Treslin/TICRR fragments were transiently transfected into 293T cells before immunoprecipitation from cell lysates using control IgG (IgG IP) or rabbit anti-MTBP (amino acids 1-284) (MTBP-IP). Lysates and precipitates were immunoblotted with detection by rat anti-MTBP (12H7) and anti-HA antibodies. VWA: von Willebrand A domain; β, β-barrel

A structural analysis between Treslin/TICRR and Ku70 confirmed the presence of a Ku70-like β-barrel domain. Initially, the Treslin/TICRR β-barrel domain appeared to lack two consecutive anti-parallel β-strands (labelled 1 and 2 in Fig S3), which contribute to the domain’s core (Walker et al 2001). The low sequence similarity between Ku70 and Treslin/TICRR in these regions and a long insertion in the Ku70 β-barrel between β-strands 2 and 3 initially confounded our automatic methods. Nevertheless, results from the RaptorX coevolution-based contact prediction method allowed the Treslin/Ku70 alignment to be extended by 50 residues N-terminally, in which the two previously-missing β-strands in Treslin/TICRR became apparent due to a strong anti-parallel co-evolution signal (Fig S3) (Jones 1999, Wang et al 2017). To investigate whether fold recognition analysis generated consistent results, we submitted this conserved region of the human Treslin/TICRR protein to the fold assignment software I-TASSER (Iterative Threading ASSEmbly Refinement) (Yang & Zhang 2015). I-TASSER consistently recognised human Ku70 (PDB-IDs: 1JEQ_A) and the Ku70-like β-barrel in the SPOC domain (PDB-ID: 1OW1) as best templates.

In summary, three independent lines of enquiry, (i) sequence conservation (HHpred), (ii) a coevolution-based contact prediction (RaptorX), and (iii) threading (I-TASSER), provided strong and consistent evidence that the conserved M domain in Treslin/TICRR folds as a Ku70- like β-barrel containing seven core β-strands (Figs 3A and S3) (Vangone et al 2011). Ku70-like β-barrel domains have been described with roles mainly related to dimerization.

### A Ku70-like β-barrel domain is also present in Sld3 and Sld7/MTBP

Using DALI structural comparison searches we also identified previously unappreciated Ku70- like β-barrels in both the Sld3 binding domain of yeast Sld7 and the Sld7-binding domain of Sld3 (PDB-ID: 3X37_B) (PDB-ID: 3X37_A) (Figs 3A and S3) (Itou et al 2015). The Sld3 β-barrel is truncated, containing only five β-strands. In summary, the Sld3/Sld7 heterodimer forms in a structurally equivalent manner to the Ku70/Ku80 heterodimer that is a homotypic dimer of two structurally similar domains.

The homology of the β-barrel domains for Sld3 and Sld7 and their respective human orthologues Treslin/TICRR and MTBP (Boos et al 2011, Boos et al 2013, Kohler et al 2019, Kumagai & Dunphy 2017, Sanchez-Pulido et al 2010) suggest that the Ku70-like β-barrel newly identified in Treslin/TICRR is an excellent candidate for being the principal region (heterodimerization domain) that interacts with MTBP.

### The Ku70-like β-barrel of Treslin/TICRR is required for interaction with MTBP

To test whether Treslin/TICRR and MTBP may indeed interact via a homotypic Ku70/Ku80- type β-barrel-dependent interaction, we characterised this interaction. Previous biochemical and structural studies had shown an involvement of MTBP/Sld7 elements, now established here as part of the β-barrel, in the interaction with Treslin/Sld3 (Itou et al 2015, Kohler et al 2019).

We showed previously that deleting two large regions of the Treslin/TICRR M domain, amino acids 265-408 (M1) or 409-593 (M2), compromised MTBP binding (Boos et al 2013). Deleting M2 abrogated and deleting M1 severely weakened this interaction. Figure 3B shows that a fragment of Treslin/TICRR containing amino acids 260-671 that included M1 and M2 co-immunoprecipitated with endogenous MTBP in lysates of transfected 293T cells. To test the involvement of the Ku70-like β-barrel in Treslin/TICRR, we deleted amino acids 370-400 and 401-420, each containing portions that aligned with Sld3 regions that entertain direct contacts with Sld7 (Fig 3A, * symbols) (Itou et al 2015). Both deletions severely compromised the interaction with MTBP (Fig 3B), indicating that the β-barrel is required.

We found that a region C-terminal to the Ku70-like β-barrel is also required for MTBP interaction. The N-terminal 557 amino acids of Treslin/TICRR, but not the N-terminal 486 amino acids, bound to MTBP (Fig 3B). Small deletions revealed that the amino acids 518-543, but not 487-517 and 545-557, are required for MTBP binding (Fig 3B). In yeast Sld3, a short sequence approximately 35 amino acids C-terminal to the β-barrel also contains six amino acids that directly contact Sld7 (Itou et al 2015). We conclude that the Ku70-like β-barrel in the Treslin/TICRR M domain cooperates with a second region further to the C-terminus in MTBP binding.

Together, our analysis of the N-terminal 600 amino acids of Treslin/TICRR revealed that the structurally conserved part with Sld3 includes the K70/80-like β-barrel in the M domain. Thus, the central part of the Treslin/TICRR protein including the M, STD and TDIN domains constitutes a core that is homologous to Sld3, flanked by Treslin/TICRR-specific terminal domains.

### The Sld3-homologous Treslin/TICRR core is insufficient to support replication

We next wanted to test whether the Sld3-like Treslin/TICRR core is sufficient to support replication in human cells or whether it requires the higher eukaryote-specific CIT and C-terminal domains. We performed BrdU-PI flow cytometry upon RNAi-replacement of Treslin/TICRR using mutants that lacked either the CIT (Treslin/TICRR-ΔCIT, amino acids 1-264 deleted), the C-terminal region (Treslin/TICRR-ΔC853, C-terminal 853 amino acids deleted), or both (Treslin/TICRR-core) (Fig. 4A). Treslin/TICRR-ΔCIT and Treslin/TICRR-ΔC853 cells showed relatively normal BrdU-PI profiles compared to Treslin/TICRR-WT cells, with S phase populations clearly separated from G1 and G2/M cells by higher BrdU signal intensities (Fig. 4B). Quantification of multiple independent experiments indicated mild reductions in Treslin/TICRR-ΔCIT and Treslin/TICRR-ΔC853 lines (Fig. 4C). Testing additional clones confirmed these results (Fig. S4A, B, D and E), although, as described for Treslin/TICRR-ΔSTD, there was some clonal variability, with one of three ΔC853 clones (no. 29) rescuing like Treslin/TICRR-WT (Fig. S4E). Expression levels of Treslin/TICRR-ΔC853 clones were similar or higher than Treslin/TICRR-WT (Fig S4B and D).

**Figure 4.**
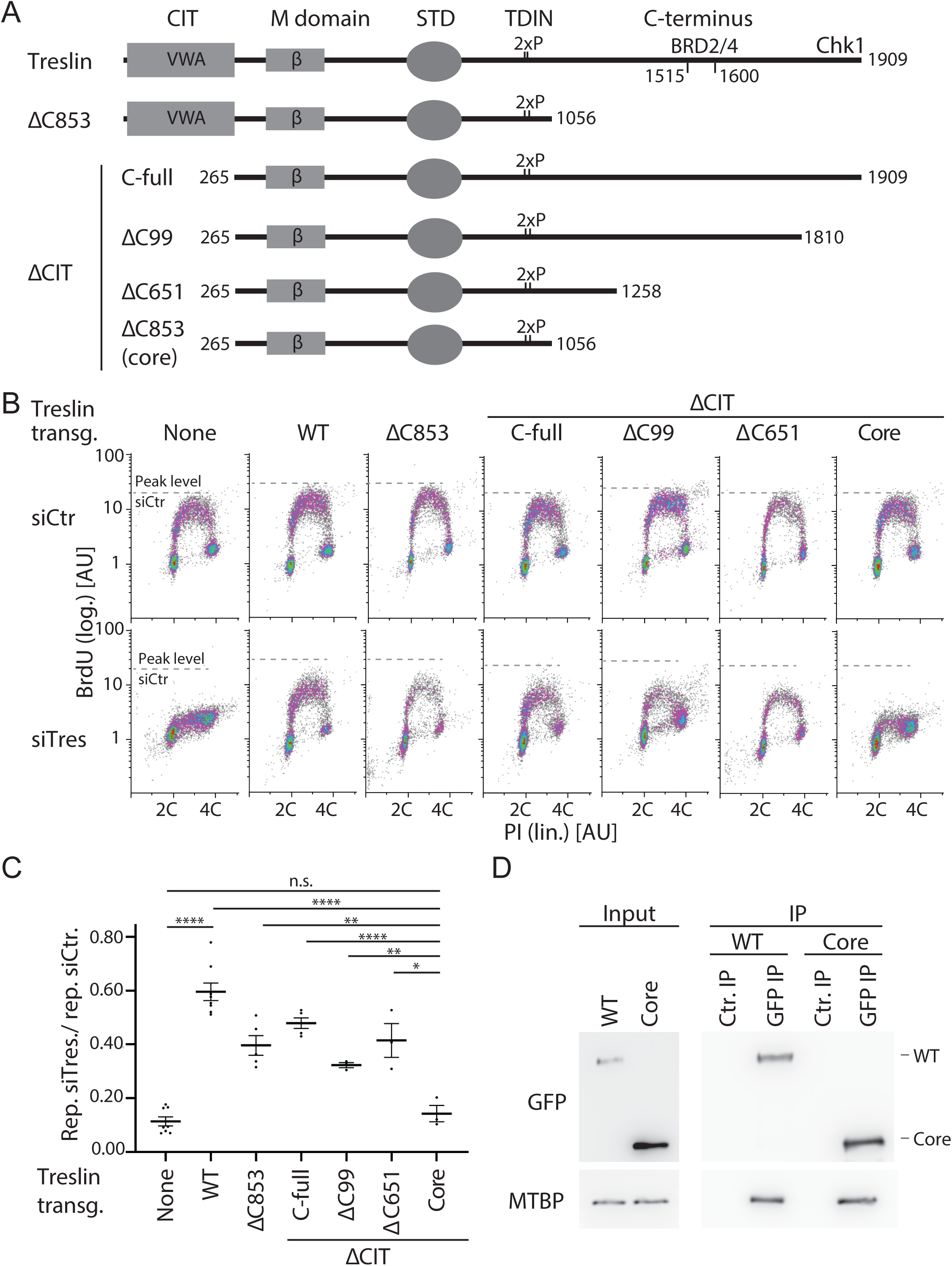
The CIT and the region between amino acids 1057-1257 of Treslin/TICRR cooperate to support replication in human cells. **(A)** Schematic representation of Treslin/TICRR mutants used in this figure. Δ: deletion; C99, 651, 853: C-terminal 99, 651 or 853 amino acids, Chk1 kinase binding requires the C-terminal 99 amino acids, BRD2/4 binds to a region between amino acids 1515 and 1600 that were deleted in Treslin/TICRR-ΔC651, -ΔC853, -ΔC394 and -ΔC309 (latter two mutants shown in Fig. S5), respectively. ΔCIT, amino acids 1-264 deleted. **(B)** Flow cytometry density plots of experiments as described in Fig. 2B using the stable U2OS cell lines expressing siTreslin-resistant Treslin/TICRR mutants described in A. Cell clones: ΔC853-5, ΔCIT(-C-full)-5; ΔCIT-ΔC99-25; ΔCIT-ΔC651-61; core-35. **(C)** Quantification of relative overall replication as described in Fig. 2C of several independent experiments as described in B). Cell clones as in B; Error bars: SEM; sample numbers (n): 8 (none; WT), 5 (ΔCIT(-C-full); ΔC853), 3 (ΔCIT-ΔC99; ΔCIT-ΔC651; core); significance tests: parametric, unpaired, two tailed student t-test, *P≤0.05. **(D)** Immunoblot with mouse anti-GFP or rat anti-MTBP (12H7) antibodies of co-immunoprecipitation experiment using 293T cells transiently transfected with GFP-Flag-Treslin/TICRR-WT or core. Native lysates were immunoprecipitated with anti-GFP nanobodies (GFP-IP) or empty control beads (Ctr. IP).

Surprisingly, the Treslin/TICRR-core mutant was inactive. BrdU incorporation in Treslin/TICRR- core cells was nearly as strongly suppressed as in the non-replicating control lines (Fig 4B and C, additional clones in Fig S5A-C). This indicated that, albeit individually non-essential for replication, simultaneous deletion of both terminal regions had an additive or even synergistic effect on DNA replication. We concluded that the Sld3-like core domains of Treslin/TICRR require the CIT domain and the C-terminal region to support replication in human cells.

### The CIT cooperates with amino acids 1057-1257 in the C-terminus to support origin firing

We then tested which part of the C-terminal region cooperates with the CIT, and whether the cooperation depends on the described binding activities for Chk1 and BRD2/4. We successively truncated the C-terminal sequence in combination with CIT deletion. Truncating neither the Chk1- (Treslin/TICRR-ΔCIT/ΔC99) (Guo et al 2015) nor the Chk1- and BRD2/4- binding domains (Treslin/TICRR-ΔCIT/ΔC651) (Sansam et al 2018) recapitulated the synergistic effect (Fig 4A-C; additional clones in Figs S5 and S6). These double-deletion mutants supported replication to a level similar to Treslin/TICRR-ΔCIT. The C-terminal truncations Treslin/TICRR-ΔC651 and ΔC99 (that contained the CIT) did not greatly affect BrdU incorporation (Figs S4C and S6B). We confirmed these results with two independent double-deletion mutants: Treslin/TICRR-ΔCIT/ΔC309 that contains the BRD2/4 binding site, and Treslin/TICRR-ΔCIT/ΔC394 that does not (Fig S6A, D and E).

Treslin-core did not support replication, as described above. To test whether the known core activities of Treslin/TICRR are intact in the Treslin/TICRR-core protein we tested association with MTBP and TopBP1. Treslin/TICRR-core and ΔC853 co-immunoprecipitated TopBP1 from 293T cell lysates similarly as Treslin/TICRR-ΔC651 (with or without CIT), suggesting that C-terminal deletion of the important amino acids 1057-1257 did not detectably compromise TopBP1 binding (Fig S7, lanes 4-7). Comparison of Treslin/TICRR-core and ΔC853 with Treslin/TICRR-full-length was difficult because of differences in expression levels and blotting efficiency in transient transfections as a result of considerable size differences. Treslin/TICRR- core also bound MTBP. Some experiments (that had the same limitations as explained for TopBP1 binding experiments) suggested slightly less MTBP bound to Treslin/TICRR-core than to Treslin/TICRR-WT (Figs 4D and S6), which could indicate that the vWA domain-containing CIT makes a small contribution to MTBP binding, similarly to the vWA domain in Ku70/Ku80 (Walker et al 2001). We cannot formally rule out that the mild reduction of MTBP binding fully explains the strong replication deficiency of Treslin/TICRR-core, although this is less likely.

We conclude that two higher eukaryote-specific Treslin/TICRR regions (specifically, CIT and the C-terminal amino acids 1057-1257) have important functions in replication.

### Treslin/TICRR-core expressing cells are defective in origin firing

Subtle particularities in cell cycle profiles of Treslin/TICRR-core cells suggested that this mutant may have other defects than cells lacking Treslin/TICRR function. For example, a delay in S phase entry in Treslin/TICRR-core cells could explain the occasionally observed decrease of the S phase sub-population (Fig S5C, clone 41). To exclude secondary effects of long-term siRNA treatment as much as possible, we tested whether Treslin/TICRR-core cells showed a specific defect in origin firing in the first S phase after replacing endogenous with transgenic Treslin/TICRR. To this end, we released Treslin/TICRR-core-expressing cells and U2OS control cells from a thymidine arrest into a nocodazole block and treated them with siRNA such that they completed S phase before siTreslin could take effect. Upon nocodazole wash-out, U2OS cells typically start replicating at around 7 h, so we chose 4 h and for 12 h to analyse BrdU-PI profiles and replisome formation. All cell lines exited from the nocodazole arrest and entered G1 phase, as indicated by 2 C DNA content at the 4 h time point (Fig 5A). As usual, a subpopulation of cells released from the arrest with a delay. Subpopulations of siCtr-treated U2OS cells and siTreslin-treated Treslin/TICRR-WT cells had started BrdU incorporation 12 h after nocodazole release. The fastest of these replicating cells had completed 30-50% of genome duplication, as judged by PI signals, showing that they had been replicating for several hours. In contrast, siTreslin-treated Treslin/TICRR-core and control cells did not replicate. To test whether Treslin/TICRR-core expressing cells have defects specifically at the origin firing step of DNA replication we analysed chromatin isolated from nocodazole-released cells. The Mcm2-7 helicase loaded normally onto chromatin in siTreslin-treated Treslin/TICRR-core G1 cells (4 h), showing that licensing was intact (Fig 5B). In contrast, replisomes did not form more efficiently with Treslin/TICRR-core than in cells without transgenic Treslin/TICRR, as indicated by PCNA and Cdc45 loading onto chromatin at 12 h in Treslin/TICRR-WT cells, but not in Treslin/TICRR-core and control cells (Fig 5B). Cyclin A blotting showed that Treslin/TICRR-core cells entered S phase normally (Fig 5C). We concluded that Treslin/TICRR-core is specifically defective in origin firing.

**Figure 5.**
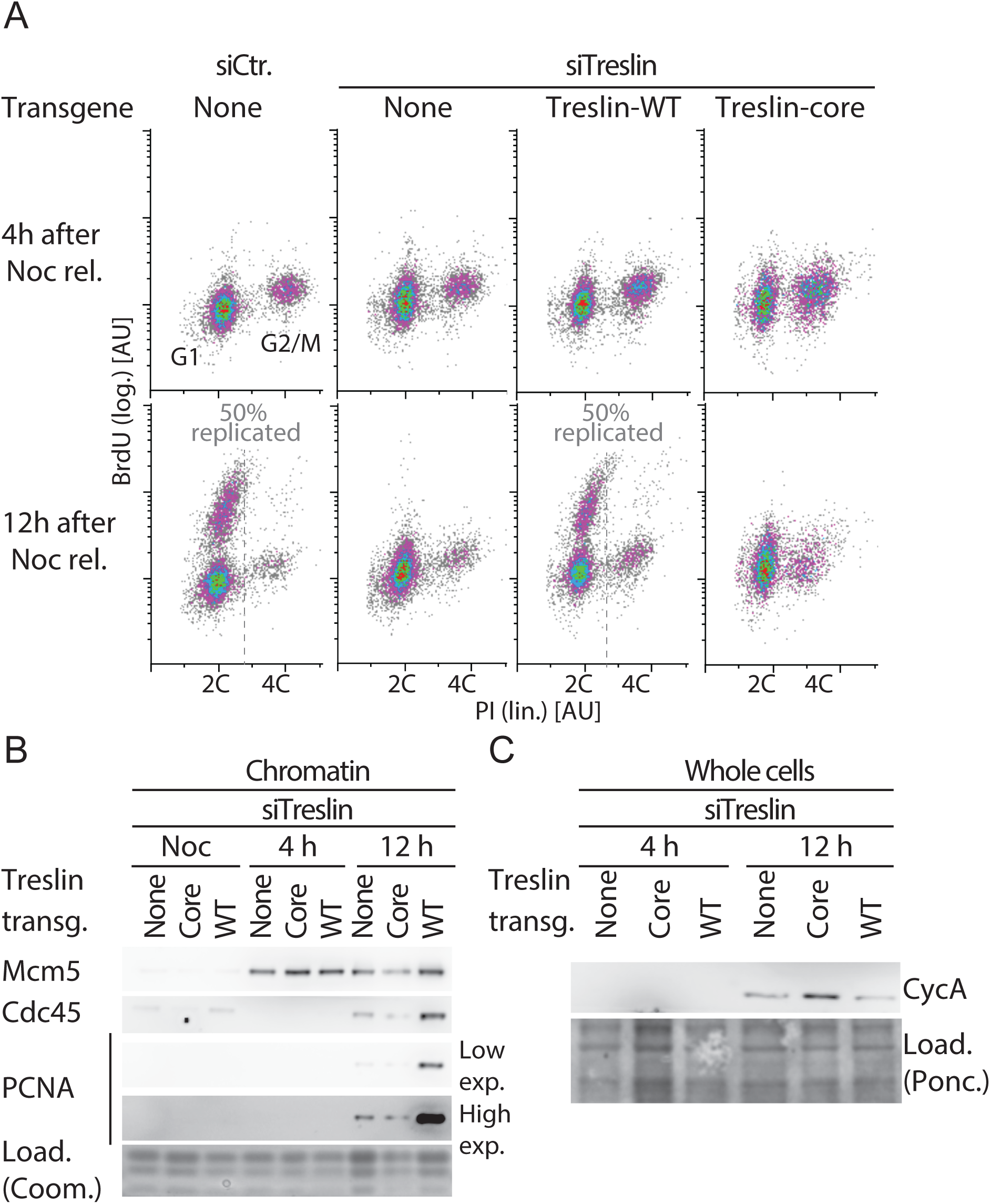
Treslin/TICRR-core does not support replisome formation. **(A)** Stable U2OS cell lines expressing no transgene or siTreslin-resistant Treslin/TICRR-WT or core were released from a thymidine arrest before treatment with siTreslin or siCtr and nocodazole. After nocodazole-release for 4 h or 12 h cells were analysed by BrdU-PI flow cytometry. Clone Treslin-core-35 was used. **(B)** Chromatin of cells treated as described in A was isolated for immunoblotting with rabbit anti-Mcm5, rat anti-Cdc45 and mouse anti-PCNA antibodies. Coomassie (Coom.) staining of low molecular weight part including histones controlled for loading. In the high exposure (exp.) the strongest band is saturated. **(C)** Whole cell lysates of cells treated as described in A were immunoblotted using mouse anti-cyclin A antibody.

Together, the Treslin/TICRR terminal regions that are specific to higher eukaryotes cooperate in parallel pathways towards an essential function in replication origin firing.

## Discussion

We here present a characterization of a major origin-firing regulator, Treslin/TICRR, based on its domain structure. Our insight that Treslin/TICRR and Sld3 share similarity of the M domain (Treslin/TICRR) and the N-terminus (Sld3), respectively, completes the view that the three central domains of Treslin/TICRR, M-domain, STD and TDIN, constitute a Sld3-like core that is flanked by two Treslin/TICRR-specific terminal regions, the CIT and the C-terminal region (Fig. 6). These terminal regions are required for Treslin/TICRR’s role in replication origin firing. Important molecular activities of the core domains are known (Fig. 6). TDIN is essential for replication in Sld3 and Treslin/TICRR through CDK-mediated interaction with Dpb11 and TopBP1, respectively (Boos et al 2011, Kumagai et al 2011, Tanaka et al 2007, Zegerman & Diffley 2007). The Sld3-STD binds Cdc45 (Itou et al 2014), an essential component of the replicative CMG helicase. Although the Cdc45-binding activity of the STD has not been investigated in Treslin/TICRR, conservation with Sld3 suggests that this biochemical activity might also be conserved (Itou et al 2014). We show here that the Treslin/TICRR-STD is required for replication origin firing in cultured human cells, confirming that it has retained important replication functions in humans. The M domain of Treslin/TICRR is also essential for replication in human cells and mediates the binding to MTBP (Boos et al 2013). Itou et al. showed that the M domain-equivalent of Sld3 constitutes a direct binding surface for Sld7 (Itou et al 2015). We reported earlier that the M domain interacting region in MTBP, approximately the N-terminal MTBP half, contains homology to the Sld3-binding N-terminus of Sld7 (Kohler et al 2019). Here we show that the interaction is mediated by Ku70-like β-barrel domains in Treslin/TICRR/Sld3 and MTBP/Sld7 (Itou et al 2015, Kohler et al 2019), suggesting that they form homotypic dimers comprised of structurally similar domains, similar to Ku70-Ku80 dimerization (Walker et al 2001). Uncharacterised important molecular activities might be situated in the regions between the domains with proven homology to Sld3, such as the DDK-dependent binding to the Mcm2-7 helicase shown for a short stretch of amino acids between the STD and TDIN of Sld3 (Deegan et al 2016).

**Figure 6.**
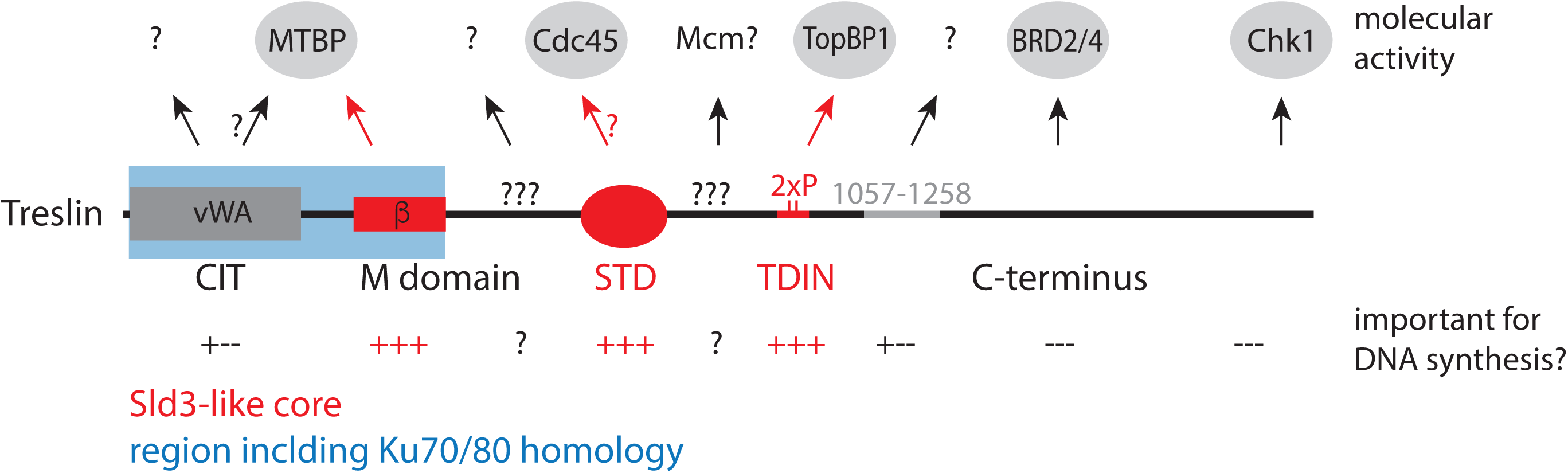
Treslin/TICRR domain structure with described molecular activities. The Sld3-like core of Treslin/TICRR is formed by the Ku70/80-like β-barrel, the STD and the TDIN domains (red). All core domains are essential for replication. Two non-core domains, the CIT domain that contains a Ku70/80-like vWA fold and the domain between amino acids 1057 and 1257, cooperate to support DNA replication in human cells. Binding regions of described interactors are shown. “?” indicates that undescribed/hypothetical activities may reside in structurally uncharacterised regions of the protein.

We found that the Sld3-like core of Treslin/TICRR was insufficient to support replication and origin firing in U2OS cells, whereas individual deletions of the Treslin/TICRR-specific CIT and C-terminus had only mild effects, if any (given the uncertainty due to clonal variability), on Treslin/TICRR’s ability to support replication. We concluded that the CIT and the C-terminal region cooperate in parallel pathways to promote DNA replication origin firing. The simplest scenario is that CIT and the C-terminal region promote firing through functions in the molecular process of origin firing that have yet to be revealed. However, more indirect scenarios cannot be excluded. Our finding supports the idea of molecular processes and regulations that are specific to higher eukaryotes to facilitate faithful duplication of their extremely complex genomes. Previous publications had shown roles for higher eukaryote-specific protein domains of TopBP1 (Kumagai et al 2010) and MTBP (Kohler et al 2019).

The molecular activities underlying the proposed origin firing functions of CIT and the C-terminal region remain unknown. Our mutants combining CIT-deletion and successive C-terminal truncation excluded significant contributions of the described Chk1- and BRD2/4- binding regions of the Treslin/TICRR C-terminus (Guo et al 2015, Sansam et al 2018). Instead, comparing the Treslin-ΔCIT/Δ853 with Treslin-ΔCIT/Δ651 mutants suggested that the relevant activity is situated between amino acids 1057 and 1257 of human Treslin/TICRR. Because this region is very close to the TDIN we considered that TopBP1 binding could be compromised in Treslin-Δ853. Although minor defects of Treslin/TICRR-Δ853 mutants in TopBP1 binding cannot be formally excluded we found no clear evidence for such a defect, regardless of whether or not the CIT was present. Also the fact that Treslin-Δ853 mutants that contain the CIT have mild or no defects in supporting genome replication, depending on the clone observed, argues against a significant TopBP1 binding deficiency. A relevant activity in the CIT for origin firing may be to support the binding to MTBP for two reasons: 1) Treslin/TICRR-core and Treslin/TICRR-ΔCIT bound somewhat less well to MTBP (Fig 4D and (Kohler et al 2019)), and 2) the CIT-equivalent domain in Ku70/80 makes a small contribution to the Ku70/80 dimer interface (Walker et al 2001). This potential mild MTBP binding defect may contribute to the inability of Treslin/TICRR-core to support origin firing. However, we find it unlikely that such a moderate defect fully explains the strong replication deficiency of Treslin/TICRR-core. This view is supported by the fact that a Sld3/Sld7-type interaction does not necessarily require a CIT, because Sld3-Sld7 dimerization is CIT-independent. We cannot formally exclude that Treslin/TICRR-core is prone to unfolding, although its normal expression levels and good TopBP1 and MTBP binding capability speak against this. Other labs also reported that C-terminally deleted Treslin/TICRR-ΔC651 supported replication well (Kumagai et al 2010), suggesting that C-terminal truncation is compatible with Treslin/TICRR’s capability to support replication.

Interestingly, the CIT contains a vWA domain that is also shared by Ku70/Ku80 (Walker et al 2001). A specific molecular activity of the CIT cannot be delineated from the presence of this domain since vWA domains in other proteins have a variety of activities. The Ku70/80 similarity in the CIT and M domains supports speculation that, during evolution, Treslin/TICRR received the CIT and the M domain in a single event of genomic recombination. The identical order of the domains in the Ku70/80 proteins suggests that Ku proteins and Treslin/TICRR share an ancestral donor for these domains or that one of the two (Ku and Treslin/TICRR) was the donor. Because animal and plant Treslins contain CITs, the last common ancestor of plants and animals likely contained a CIT. As opisthokonts, fungi and animals are more closely related to each other than animals are to plants, so the CIT must have been lost from Sld3 during yeast evolution. In conclusion, the CIT may have been “donated” to Treslin/TICRR as one unit alongside the Ku70-like β-barrel. Both together had the capability to form homotypic dimers with MTBP. The minor (or absent) contribution of the CIT to MTBP binding presents the possibility that it was retained in most branches of evolution due to another function important for eukaryotic cells.

Determining the molecular and cellular functions of the non-core Treslin/TICRR domains will help us better understand the specifics of origin firing in higher eukaryotes compared to yeast. Because Treslin/TICRR mediates origin firing regulation, understanding its non-core domains will likely be necessary to unravel how the complex higher eukaryotic cells coordinate origin firing with other cellular processes.

## Materials and Methods

### Cell culture

U2OS (ATCC-HTB-96) and 293T (ATCC CRL-11268) cells (both lines kind gift from The Crick institute tissue culture) were cultured in standard conditions in DMEM/high glucose (Life Technologies, 41965062), 10 % FCS, Penicillin/Streptomycin in 5 % CO_2_. Stable Treslin/TICRR- expressing U2OS cell clones were generated using a pIRES puro3-based vector system by random genome integration followed by selection on 0.3 µg/ml puromycin and picking of individual clones as described (Boos et al 2011, Boos et al 2013).

### Analysis of unsynchronised and synchronous stable U2OS cells by BrdU-flow cytometry and chromatin analysis

Endogenous Treslin/TICRR was replaced by siTreslin-resistant transgenes by transfecting U2OS cells twice with Treslin/TICRR siRNA (GAACAAAGGTTATCACAAA) using RNAiMax (Life Techmologies/ 13778150) as described (Boos et al 2011). Luciferase siRNA (GL2, Dharmacon) served as a control. For end point analysis of unsynchronised cells, cells were labelled with 10 µM BrdU for 30 min 72 h after the first transfection, harvested and stained with anti-BrdU-FITC (Becton Dickinson/ 556028) and propidium iodide as described (Boos et al 2011). Flow cytometry analysis was performed, analyzed and quantified as described (Kohler et al 2019). In brief, for quantification of replication rescue using BrdU-PI flow cytometry, the BrdU signal intensity of the S phase cell population was background-subtracted using the combined BrdU-channel signal of G1 and G2/M populations to determine the replication-dependent BrdU signal. This replication signal was normalized to the replication signal of siCtr-treated cells of the same cell clone to calculate the relative replication rescue. For analysis of synchronized U2OS cells in Fig 2D and E cells were arrested by treatments with 2 mM thymidine for 18 h, release for 10 h, and arrested once again with 2 mM thymidine for 18 h. 4 h after release from the second thymidine block cells were treated with siRNA and 100 µg/ml nocodazole was added for 16h. Release from the nocodazole arrest was done by washing the cells twice. After cultivation for four or twelve hours, cells were harvested and analysed by BrdU-flow cytometry as described above or by immunoblotting of whole cell lysates or chromatin-enriched fractions as described (Boos et al 2013). For Fig 5, cells were instead treated with siRNA and arrested by treatment with 2 mM thymidine for 20 h. Upon release from the thymidine block, 100 µg/ml nocodazole was added for 18h. Cells were treated with the second round of siRNA 4 h after the start of the nocodazole arrest.

### Antibodies and affinity matrices

Antibodies against Treslin, MTBP and TopBP1 were described (Boos et al 2011, Boos et al 2013, Kohler et al 2019). Anti-BrdU-FITC (Becton Dickinson/ 556028); anti-HA (mouse, 16B12; Covance); anti-GFP nanobodies (kind gift from Kirill Alexandrov); anti-GFP (mouse, JL-8, Clonetech, 632381), anti-Mcm5 (rabbit, ab17967, abcam), anti-Cdc45 (rat, 3G10, kind gift from Helmut Pospiech), anti-PCNA (mouse, sc-56, Santa Cruz), NHS sepharose (Fisher Scientific, 10343240), Protein G magnetic beads (Life Technologies, 10004D)

### Immunoprecipiation from transiently transfected 293T cell lysates

293T cells were transfected using standard calcium phosphate precipitation. 72 h after transfection, cells were harvested and lysed in 5-10 times cell pellet volume using detergent in native lysis buffers and douncing. Lysis buffer for anti-GFP immunoprecipitations in Fig S7 was 20mM Hepes, 250mM NaCl, 10% Glycerol, 0,1%Triton, 2mM EDTA, 10mM NaF, 2mM mM ß-Mecaptoethanol, Complete EDTA-free protease inhibitors (Roche, 5056489001); for Fig 4D lysis buffer was 20mM Hepes, 300mM NaCl, 10% Glycerol, 0,1%Triton, 2mM EDTA, 2mM mM ß-Mecaptoethanol, Complete EDTA-free protease inhibitors (Roche, 5056489001); for rabbit anti-MTBP immunoprecipitation in Fig 3B 20mM Hepes, 200mM NaCl, 10% Glycerol, 0,1%Triton, 2mM mM ß-Mecaptoethanol, Complete EDTA-free protease inhibitors. Lysates from cells from 12.5 % (Fig 3B) and 100 % (Figs 4D, S1 S7) confluent 10 cm dish (Figs S1 and S7), as well as 10 µl (Figs S1, S7 and 4D) GFP nanobody NHS sepharose beads (1 µg/µl) or 1 µg anti-MTBP (amino acids 1-284) antibody on 10 µl magnetic protein G slurry beads (Fig 3B) were used per reaction. After washing three time with lysis buffer beads were boiled in Laemmli loading buffer and analysed by SDS PAGE and immunoblotting. For CDK treatment of lysates, 67 µg/ml bacterially purified Cdk2-cyclin A (purification system generously donated by Tim Hunt), 5 mM ATP and 5 mM MgCl_2_ were added to the lysis buffers.

## Data availability

The authors will comply with Nature Research policies for the sharing of research materials and data.

## Acknowledgements

We would like to thank the members of the S Westermann, H Meyer and D Boos labs for discussion and sharing expertise and regents.

## Author contributions

Conception and design: D.B., P.F.; Development of methodology: P.F., L.S.P. D.B.; Acquisition of data: P.F., L.S.P, A.M., D.B.; Analysis and interpretation of data: P.F., L.S.P, A.M., C.P.P., D.B.; Writing and reviewing of the manuscript: D.B. (writing), L.S.P., C.P.P.; Study supervision: D.B.

## Supplementary Information

### Supplementary figure legends

**Supplementary figure S1.**
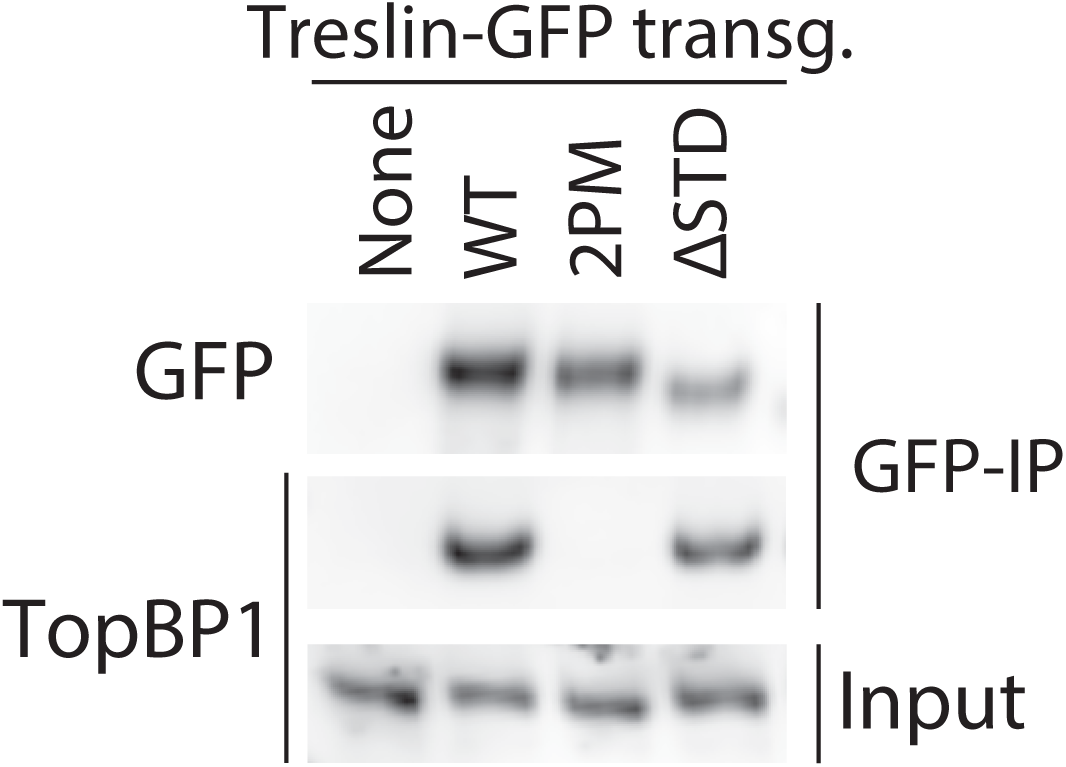
Treslin/TICRR-ΔSTD is proficient in binding TopBP1. GFP-Flag-Treslin/TICRR-WT, 2PM or ΔSTD were transiently transfected into 293T cells. Native lysates were used for anti-GFP nanobody immunoprecipitation (IP) in the presence of recombinant Cdk2-cyclin A to promote interaction with TopBP1. Lysates and bead-bound material were analysed by immunoblotting using mouse anti-GFP and rabbit anti-antibodies. Treslin/TICRR-2PM did not bind TopBP1, as expected because the relevant CDK sites in the TDIN are mutated to alanine. Treslin/TICRR- ΔSTD was able to bind to TopBP1.

**Supplementary figure S2.**
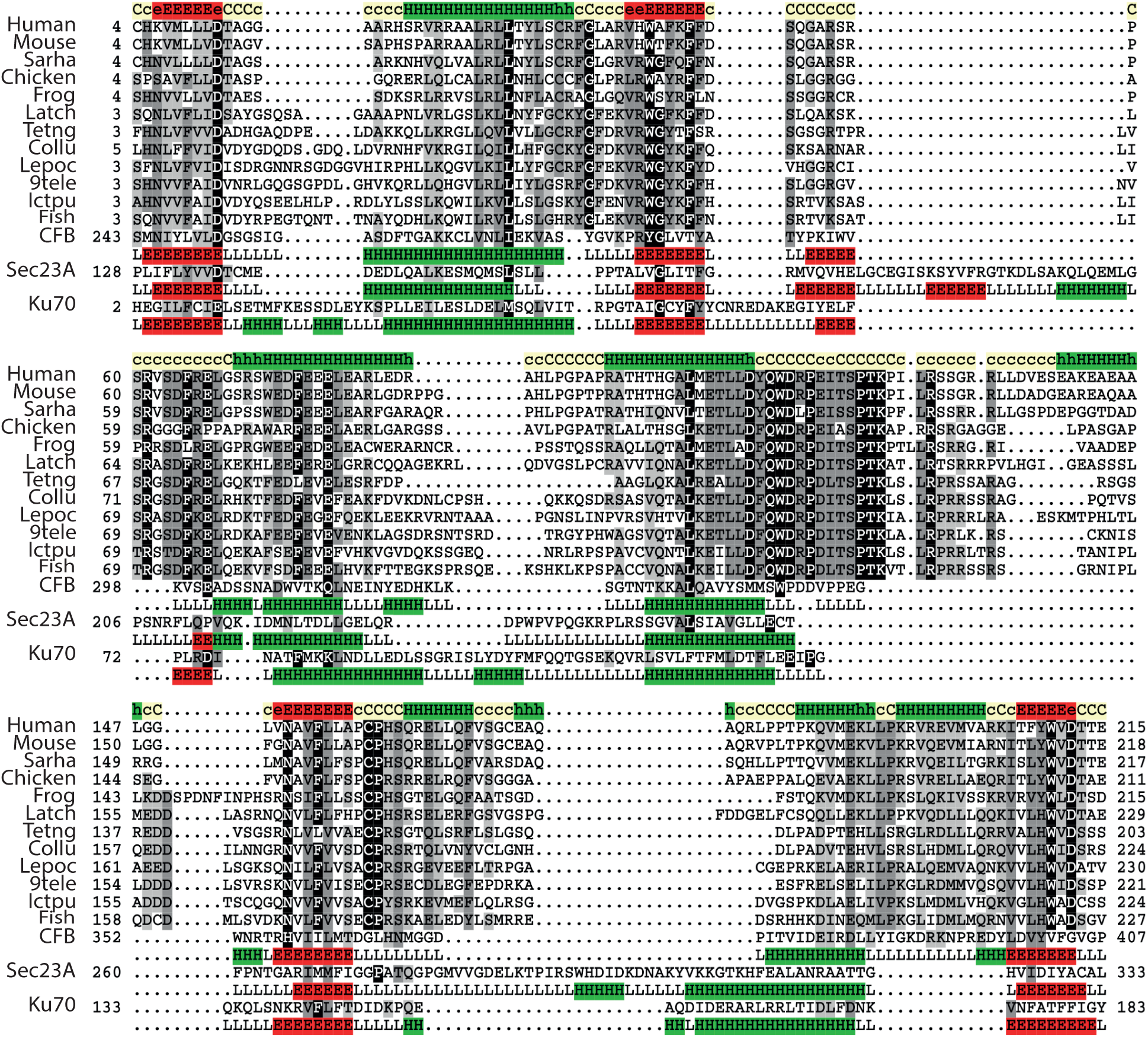
The CIT domain of Treslin/TICRR contains a vWA fold. Representative multiple sequence alignment of VWA domain in Treslin family. Secondary structure prediction using PsiPred was performed for the Treslin family, shown in the first lane; this prediction is consistent with the secondary structure of VWA domains, shown below each of the selected proteins with known structure (CFB, PDB:3HRZD; Sec23, PDB:2NUTA; Ku70, PDB:5Y58E). For figure methods and abbreviations see Figure 2A legend.

**Supplementary figure S3.**
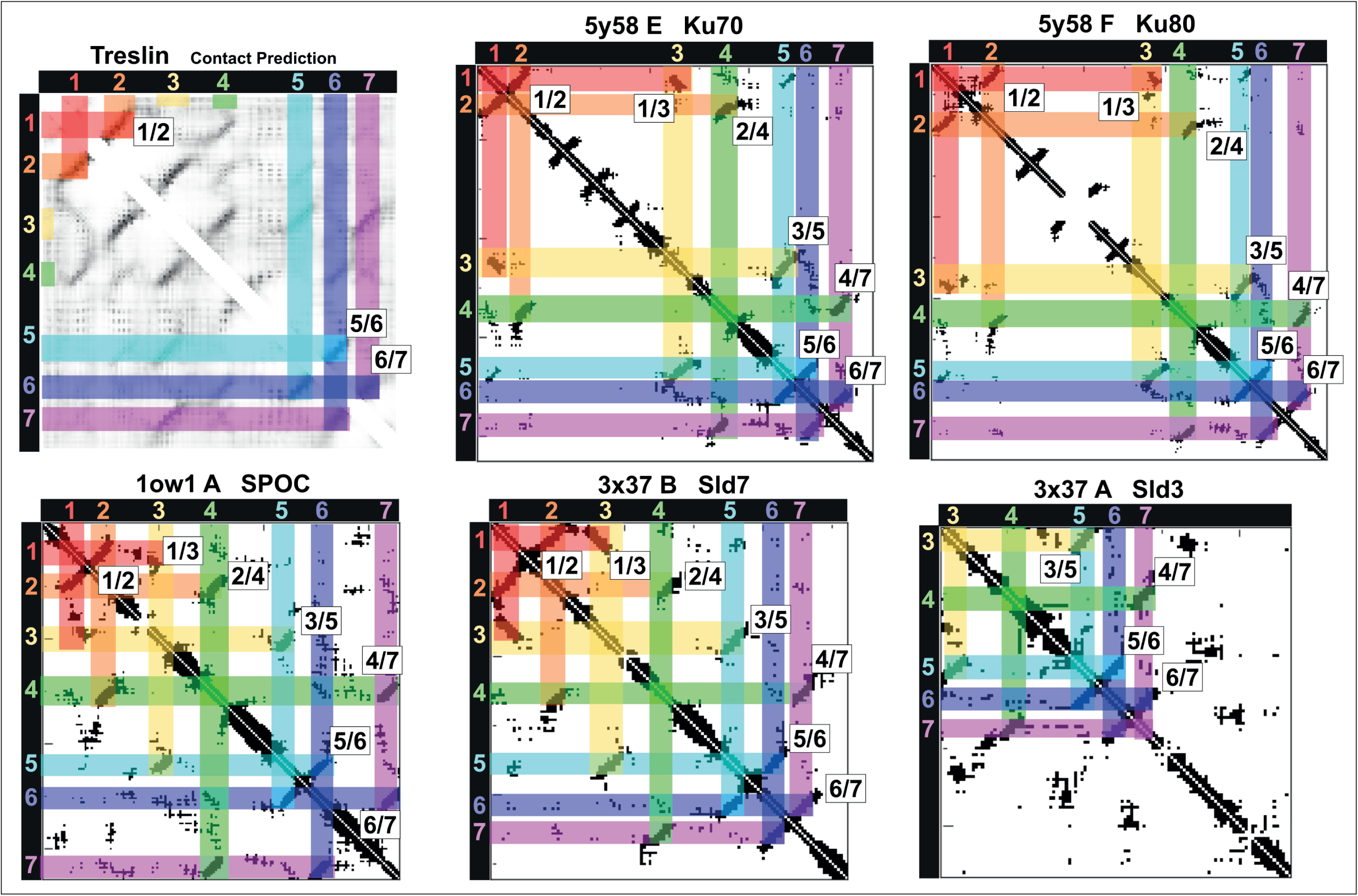
Coevolution-based and known contact maps of Ku70-like β barrel domains. RaptorX coevolution-based contact prediction for Treslin (Q7Z2Z1_HUMAN, residues 290 to 460) and known contact maps for Ku70, Ku80, SPOC, Sld7, and Sld3. Contact maps of Ku70- like β barrel domains whose structures are known were generated using the Cocomaps server (cut-off distance value = 7 Ångstroms) (Vangone et al 2011) are shown. Βeta-strands are labelled 1 to 7 and coloured in red, orange, yellow, green, cyan, violet, and purple, respectively. Β-strand contact pairs are labelled. Structural comparison with DALI using as input the Ku70 β-barrel domain core found a diverse set of proteins (Holm & Laakso 2016). This includes yeast Ku80 and human Ku70/Ku80 orthologs (XRCC5 and XRCC6) (Chen et al 2018, Nemoz et al 2018, Walker et al 2001). Other known similar structure were identified (DALI Z-Score > 7.5): the human transcriptional corepressor SHARP SPOC domain (PDB-ID: 1OW1_A) (Ariyoshi & Schwabe 2003), fission yeast Chp1 SPOC domain (PDB-ID: 3TIX_B) (Schalch et al 2011), plant FPA SPOC domain (PDB-ID: 5KXF_A) (Zhang et al 2016), and the activator interaction domain (ACID) of the human Med25 subunit of the Mediator complex (Eletsky et al 2011, Milbradt et al 2011, Vojnic et al 2011). Med25 ACID domains were recently characterised in interaction with unfolded acidic transactivation domains (TADs) (Landrieu et al 2015, Lee et al 2018) and are able to have allosteric communication between opposite β-barrel’s binding surfaces (Henderson et al 2018). Therefore Ku70-like β-barrel domains have been described with roles related to dimerisation, protein-protein and protein-DNA interaction.

**Supplementary figure S4.**
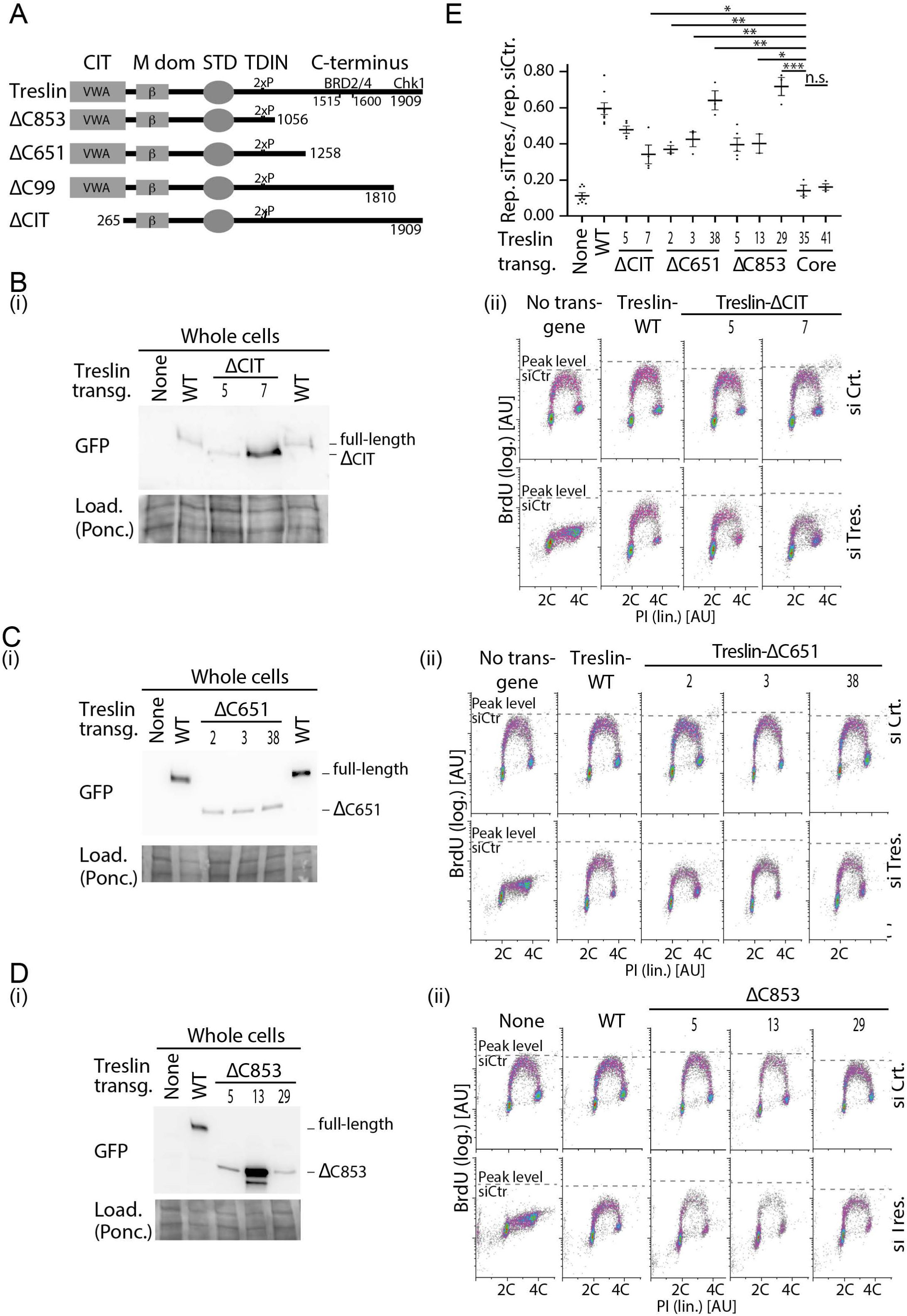
Analysis of several stable U2OS clones expressing Treslin/TICRR- ΔCIT or various C-terminal truncation mutants. A) Schematic giving an overview over the Treslin/TICRR mutants used in this figure. B-D) Immunoblots (i) to assess transgene expression levels and BrdU-flow cytometry (ii) to determine overall DNA replication of the indicated Treslin/TICRR mutants shown in A. The following U2OS clones were used for main Fig. 4: Treslin/TICRR-ΔCIT-5, ΔC853-5. Immunoblots of whole cell lysates used mouse anti-GFP and Ponceau staining (as a loading control). Flow cytometry was done after replacing endogenous Treslin/TICRR against the indicated siRNA-resistant transgenes using RNAi. Density plots are shown. Parental U2OS cells and a line expressing Treslin/TICRR-WT served to control the experiment. Dashed lines show BrdU peak level of the respective control siRNA-treated cell line in the same experiment. Clones were picked that expressed the Treslin/TICRR transgenes at similar or higher levels than Treslin/TICRR-WT to avoid under-estimating the capability of the mutants to support replication. For Treslin/TICRR-ΔC651, only low-expressing clones were found. The results are still conclusive, though, because all clones were capable to support replication. E) Quantification of overall replication in mutant Treslin/TICRR U2OS cell lines described in A-D, based on BrdU-PI flow cytometry experiments as described in B-D. For comparison, the shortest C-terminal deletion mutant (Δ99) is shown in addition to the usual control lines. The quantifications indicate that all mutants were active. It also shows the clonal variability that did not clearly correlate with expression levels, as indicated by the Treslin/TICRR-ΔC651 clones 1-3.

**Supplementary figure S5.**
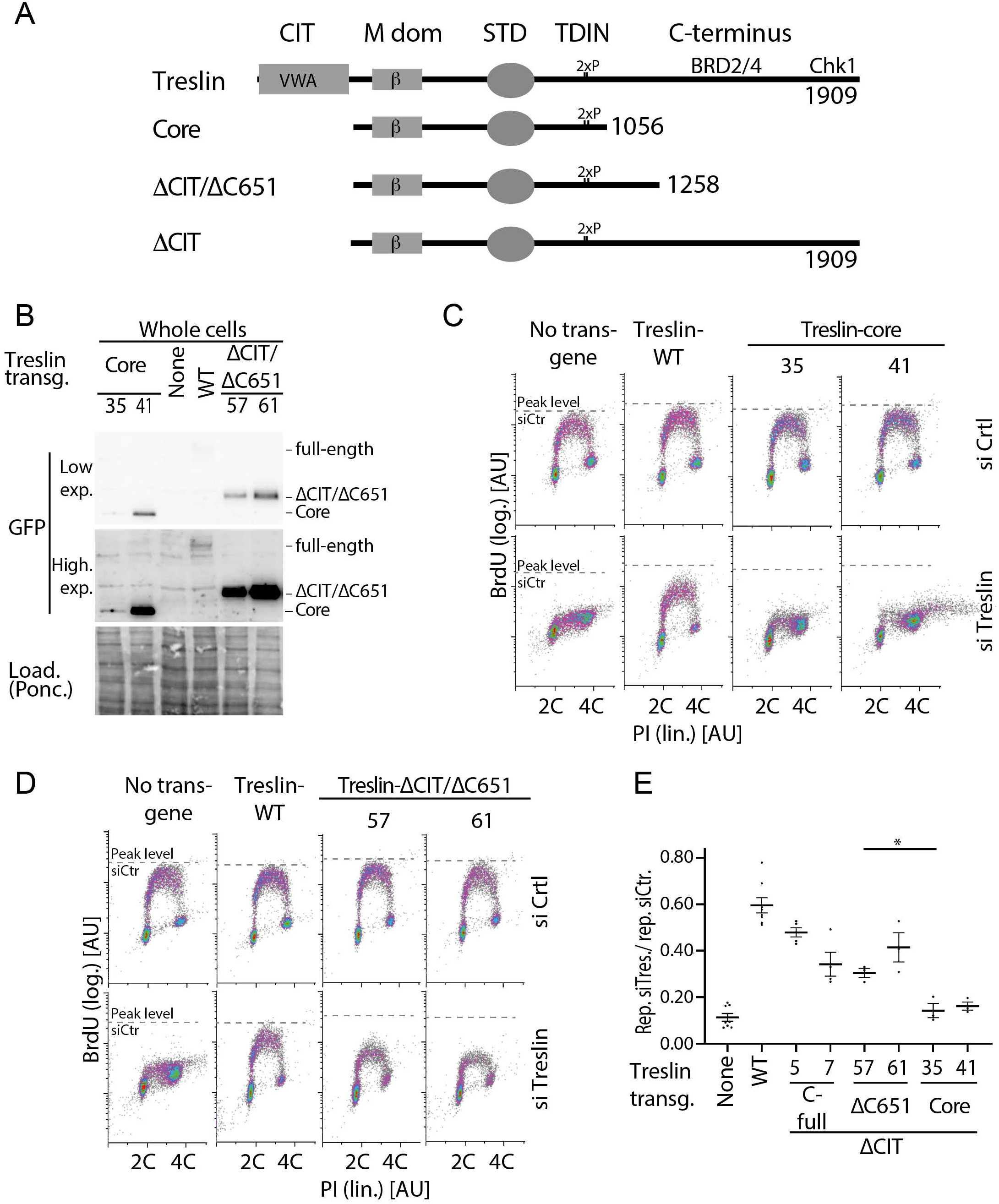
Analysis of several stable U2OS clones expressing Treslin/TICRR- core and Treslin/TICRR-ΔCIT/ΔC651. A) Schematic giving an overview over the Treslin/TICRR mutants used in B-E B-D) Immunoblots (i) and BrdU-Flow cytometry (ii) of stable U2OS cell lines expressing indicated Treslin/TICRR-mutants shown in A. Immunoblots of whole cell lysates using mouse anti-GFP and Ponceau staining (as a loading control) served to assess transgene expression levels relative to each other and Treslin/TICRR-WT. The following U2OS clones were used for main figures: Treslin/TICRR-ΔCIT/ΔC651-61 (Fig. 4), core-35 (Fig. 4, 5). Clones were picked that expressed the Treslin/TICRR transgenes at similar or higher levels than Treslin/TICRR-WT to avoid under-estimating the capability of the mutants to support replication. For BrdU-flow cytometry, density plots show overall DNA replication of stable U2OS clones shown in A. Flow cytometry was done after replacing endogenous Treslin/TICRR against the indicated siRNA-resistant transgenes using RNAi. Parental U2OS cells and a line expressing Treslin/TICRR-WT served to control the experiment. Dashed lines show BrdU peak level of the respective control siRNA-treated cell line in the same experiment. E) Quantification of overall replication in mutant Treslin/TICRR U2OS cell lines described in A, based on BrdU-PI flow cytometry experiments as described in B. For comparison, Treslin/TICRR-ΔCIT containing the full C-terminus and Treslin/TICRR-core are shown in addition to the usual control lines. Treslin/TICRR-ΔCIT/ΔC651 supports replication to levels comparable with Treslin/TICRR-ΔCIT. The exact level of replication depended on the clone used. No Treslin/TICRR-ΔCIT/ΔC651 clone, however, supported replication as poorly as Treslin/TICRR-core that showed replication similar to control U2OS cells not expressing a siRNA-resistant transgene.

**Supplementary figure S6.**
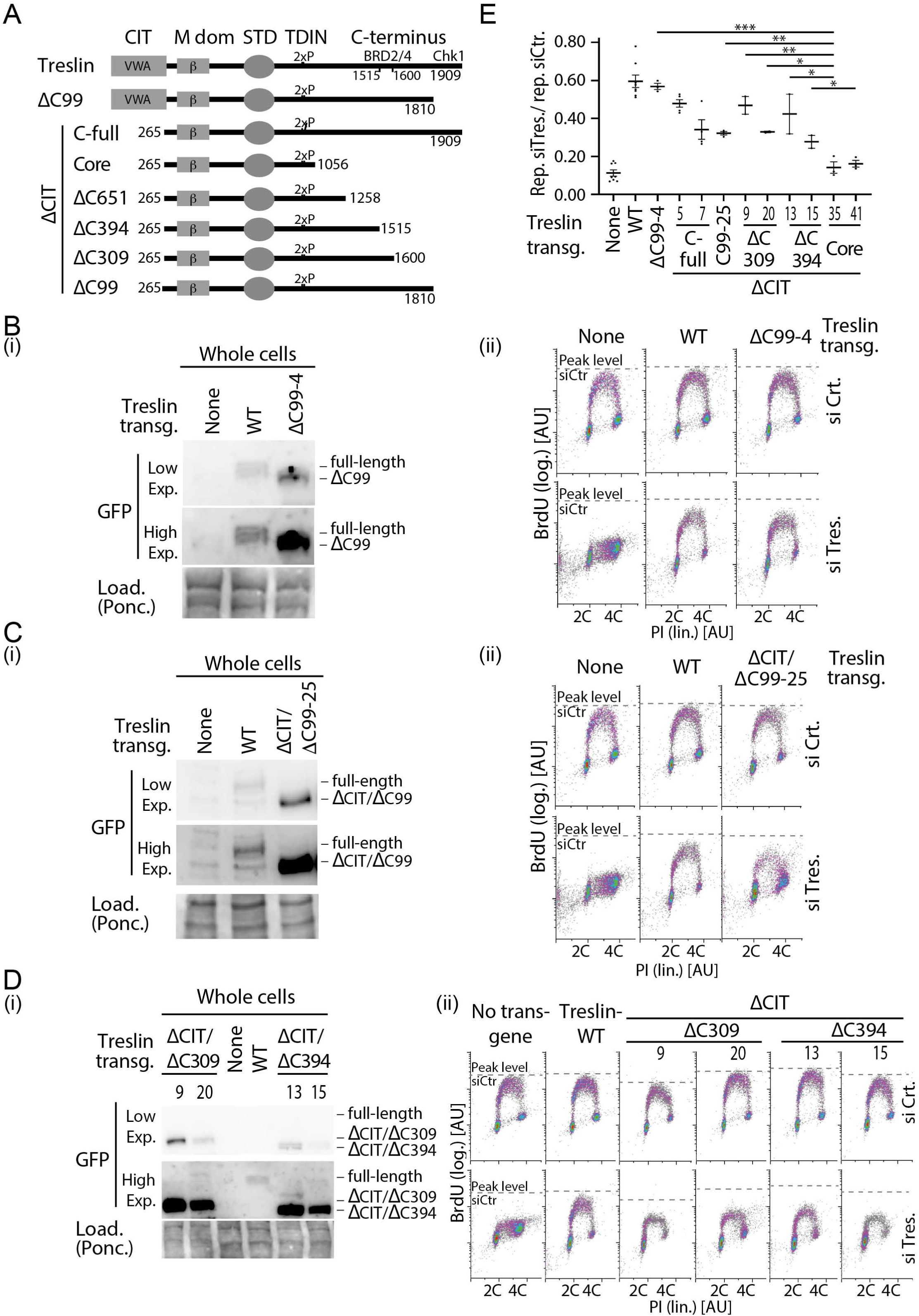
Analysis of several stable U2OS clones expressing various C-terminal truncations in combination with deletion of the CIT or Treslin/TICRR-ΔC99. A) Schematic giving an overview over the Treslin/TICRR mutants used in B-E B) Immunoblots to assess transgene expression levels of stable U2OS clones expressing Treslin/TICRR-ΔCIT/ΔC651 and ΔCIT/ΔC853. Immunoblots of whole cell lysates used mouse anti-GFP and Ponceau staining (as a loading control). Clones were picked that expressed the Treslin/TICRR transgenes at similar or higher levels than Treslin/TICRR-WT to avoid under-estimating the capability of the mutants to support replication. C/D) Density plots of BrdU-flow cytometry to determine overall DNA replication of stable U2OS clones described in A and B. Flow cytometry was done after replacing endogenous Treslin/TICRR against the indicated siRNA-resistant transgenes using RNAi. Parental U2OS cells and a line expressing Treslin/TICRR-WT served to control the experiment. Dashed lines show BrdU peak level of the respective control siRNA-treated cell line in the same experiment. E) Quantification of overall replication in mutant Treslin/TICRR U2OS cell lines described in A, based on BrdU-PI flow cytometry experiments as described in B. For comparison, Treslin/TICRR-ΔCIT containing the full C-terminus, Treslin/TICRR-ΔCIT/Δ651 and Treslin/TICRR-core are shown in addition to the usual control lines. Treslin/TICRR-ΔC99 supports replication to similar levels as Treslin/TICRR-WT. Treslin/TICRR-ΔCIT/ΔC99 supports replication to levels comparable with Treslin/TICRR-ΔCIT and ΔCIT/ΔC651, but much better than Treslin/TICRR-core. Also here, the exact level of replication depended on the clone used.

**Supplementary figure S7.**
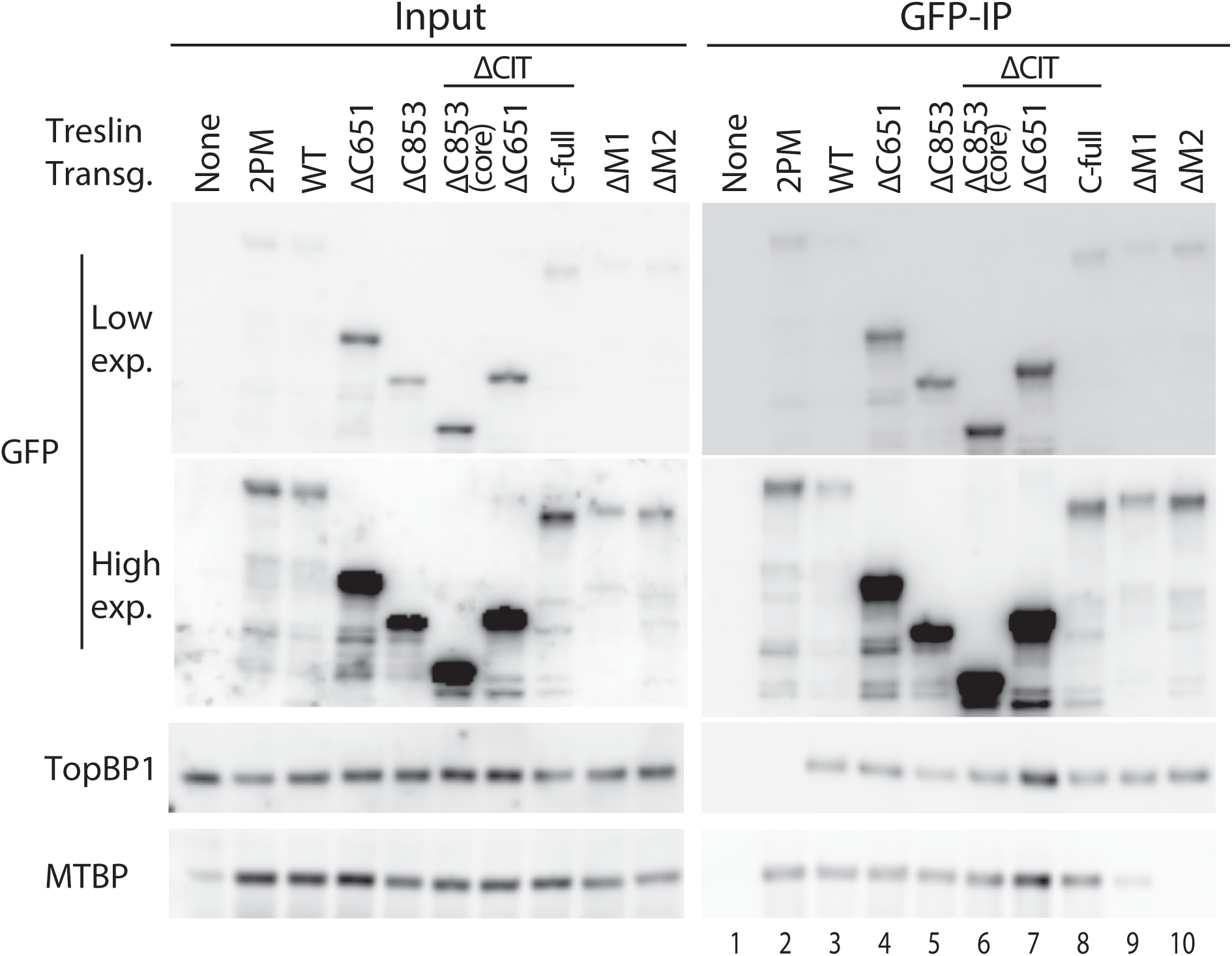
Treslin/TICRR-core is proficient in binding MTBP and TopBP1. The indicated GFP-Flag-Treslin/TICRR mutants were transiently transfected into 293T cells together with MTBP. Native lysates were used for anti-GFP nanobody immunoprecipitation (IP) in the presence of recombinant Cdk2-cyclin A to promote interaction with TopBP1. Lysates and bead-bound material were analysed by immunoblotting using mouse anti-GFP, rabbit anti-TopBP1 and rat anti-MTBP antibodies. Controls for IP specificity were made: Treslin/TICRR-ΔM1 and ΔM2 show decreased (M1) or absent (M2) MTBP signals, as expected. Treslin/TICRR-2PM did not bind TopBP1, as expected because the relevant CDK sites in the TDIN are mutated to alanine. IP capabilities using (near) full-length Treslin/TICRR versions are hard to compare by immunoblotting with those containing larger deletions because of the often weak blotting efficiency of the 210 kD full-length Treslin/TICRR. However, the smaller C-terminal truncations are better comparable. Treslin/TICRR-ΔC853 and Δ651 bound similar amounts of TopBP1 and MTBP, whether they contained CIT or not. In some experiments, however, deletion of the CIT seemed to have a minor effect on the amount of MTBP bound (Fig 4D).

